# *Allele-specific* knockouts reveal a role for *apontic-like* in the evolutionary loss of larval pigmentation in the domesticated silkworm, *Bombyx mori*

**DOI:** 10.1101/2022.05.07.490996

**Authors:** Kenta Tomihara, Peter Andolfatto, Takashi Kiuchi

**Author notes:** Corresponding Author: Kenta Tomihara, Takashi Kiuchi.

## Abstract

The domesticated silkworm, *Bombyx mori*, and its wild progenitor, *B. mandarina*, are extensively studied as a model case of the evolutionary process of domestication. A conspicuous difference between these species is the dramatic reduction in pigmentation in both larval and adult *B. mori*. Here we evaluate the efficiency of CRISPR/Cas9-targeted knockouts of pigment-related genes as a tool to understand their potential contributions to domestication-associated pigmentation loss in *B. mori*. To demonstrate the efficacy of targeted knockouts in *B. mandarina*, we generated a homozygous CRISPR/Cas9-targeted knockout of *yellow-y*. In *yellow-y* knockout mutants, black body color became lighter throughout the larval, pupal and adult stages, confirming a role for this gene in pigment formation. Further, we performed *allele-specific* CRISPR/Cas9-targeted knockouts of the pigment-related transcription factor, *apontic-like* (*apt-like*) in *B. mori* × *B. mandarina* F_1_ hybrid individuals. Knockout of the *B. mandarina* allele of *apt-like* in F_1_ embryos results in depigmented patches on the dorsal integument of larvae, whereas corresponding knockouts of the *B. mori* allele consistently exhibit normal F_1_ larval pigmentation. These results demonstrate a contribution of *apt-like* to the evolution of reduced pigmentation in *B. mori*. Together, our results demonstrate the feasibility of CRISPR/Cas9-targeted knockouts as a tool for understanding the genetic basis of traits associated with *B. mori* domestication.

**Brief abstract:** *Bombyx mori* and its wild progenitor are an important model for the study of phenotypic evolution associated with domestication. As proof-of-principle, we used CRISPR/Cas9 to generate targeted knockouts of two pigmentation-related genes. By generating a homozygous knockout of *yellow-y* in *B. mandarina*, we confirmed this gene”s role in pigment formation. Further, by generating *allele-specific* knockouts of *apontic-like* (*apt-like*) in *B. mori* × *B. mandarina* F_1_ hybrids, we establish that evolution of *apt-like* contributed to reduced pigmentation during *B. mori* domestication.

**Graphical TOC/Abstract:** 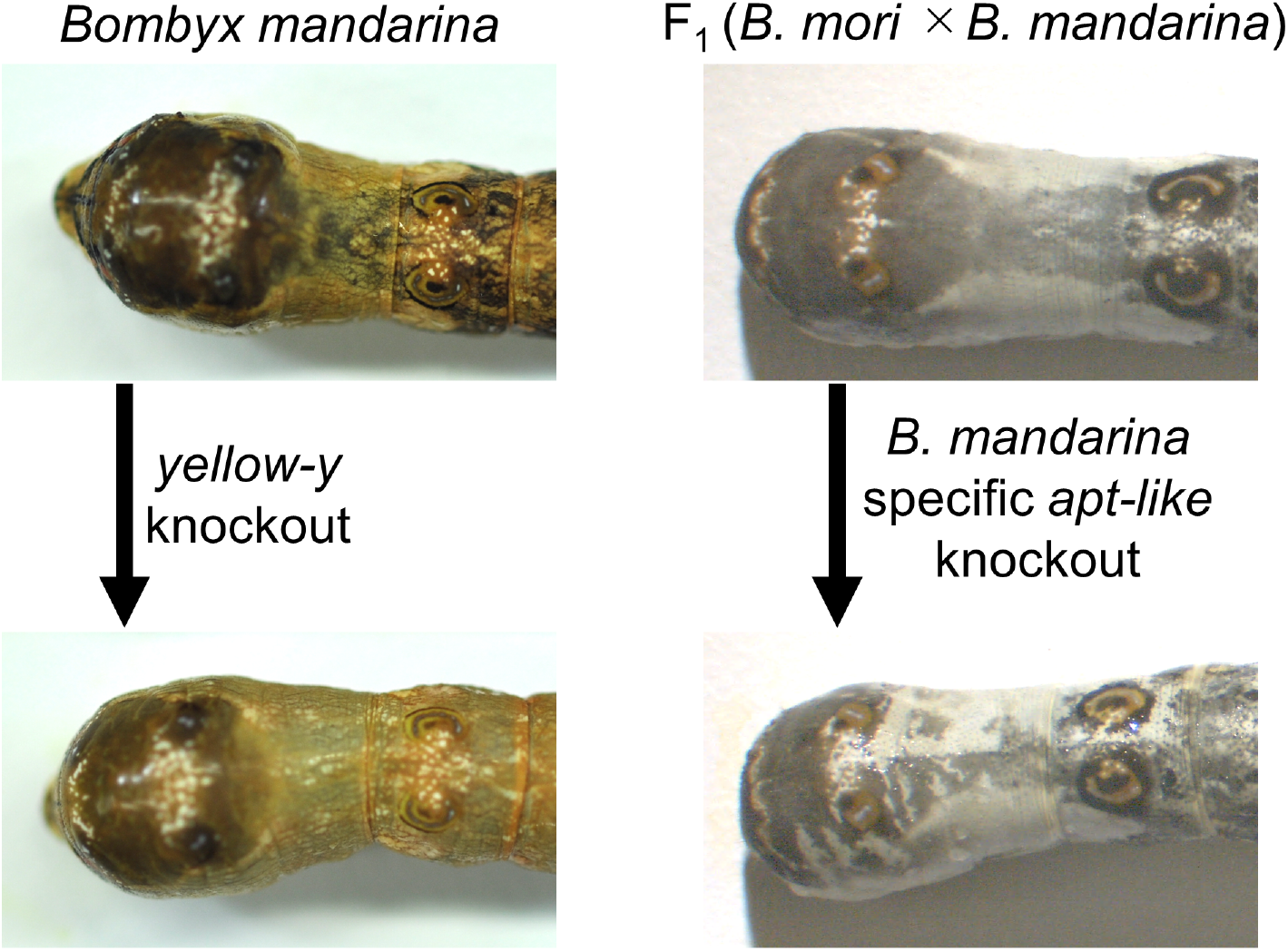

## Introduction

The silkworm, *Bombyx mori*, was domesticated over 5000 years ago from its wild progenitor species, *B. mandarina*. Under long-term artificial selection, *B. mori* acquired various characteristics suitable for sericulture. For example, the weight of the cocoon shell of *B. mori* is much higher than that of *B. mandarina* (Ômura 1950, Li *et al*. 2017, Fang *et al*. 2020). In addition, *B. mori* moths lost their flying ability due to the degeneration of their flight muscles and a reduction in wing stiffness (Lu *et al*. 2020). Among the most conspicuous domestication-associated traits is a marked reduction in pigmentation in *B. mori* larvae and adults relative to *B. mandarina* (Figure 1A). Curiously, depigmentation is a major trait contributing to the so-called “domestication syndrome” observed in a variety of domesticated animals, but for reasons that are not well-understood (Wilkins *et al*. 2014).

**Figure 1.**
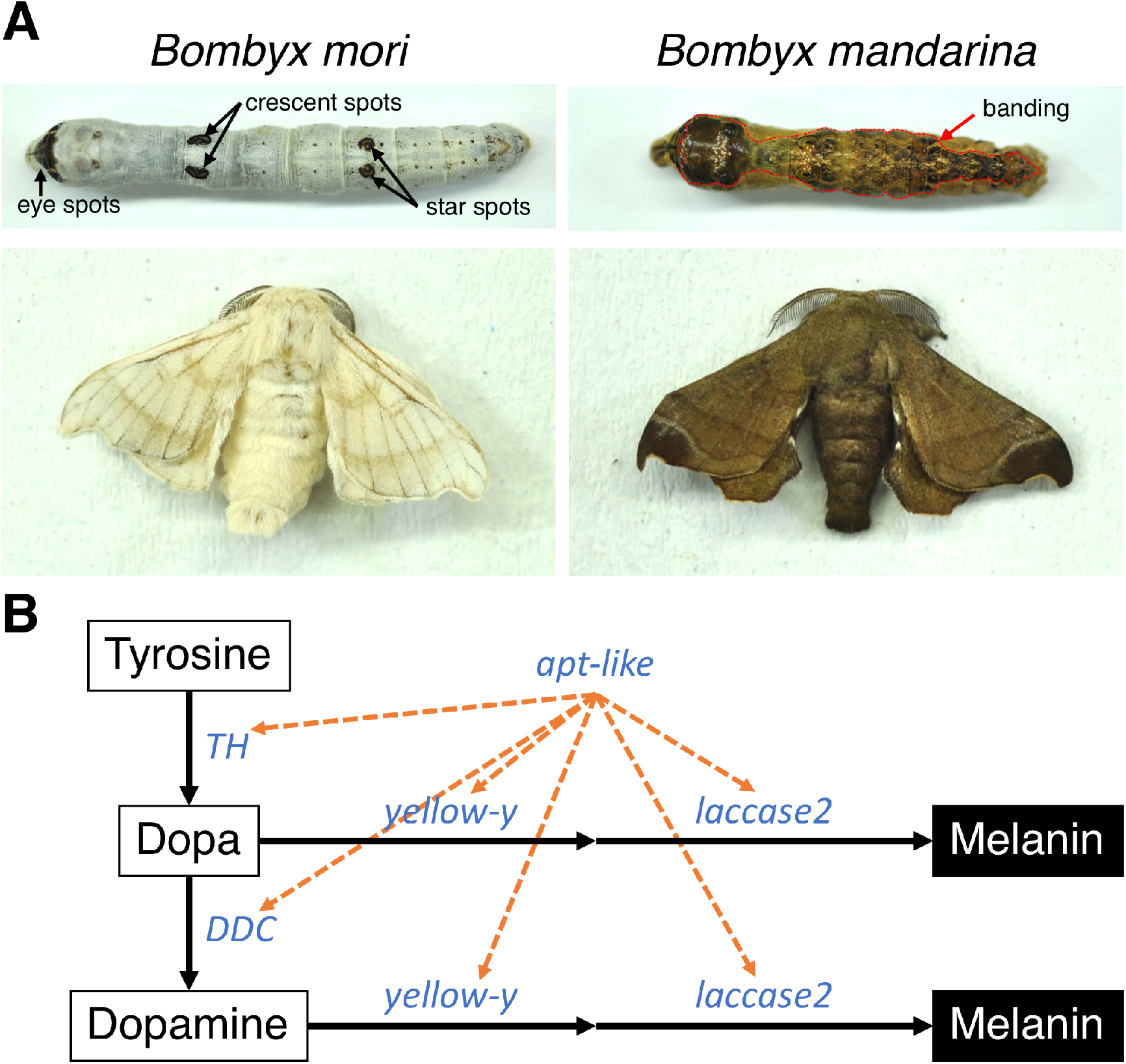
(A) Larvae (top) and adult males (bottom) of *B. mori* (left) and *B. mandarina* (right). Black arrows indicate larval spot markings (eye spots, crescent spots and star spots) and red dotted lines delimit the larval dorsal pigment patterning (“banding”). The names of spots follow the nomenclature of Yoda *et al*. (2014). (B) The proposed melanin biosynthesis pathway in *Bombyx* larvae largely adapted from Futahashi *et al*. (2008) and Yoda *et al*. (2014). Orange dashed arrows indicate presumed regulation of genes by the transcription factor *apt-like*.

The genetic basis of pigmentation loss associated with *B. mori* domestication is not yet known. Previous studies reported that melanin synthesis pathway genes *tyrosine hydroxylase* (*TH*) and *aspartate decarboxylase* (*ADC*, also known as *black*) were potentially targets of selection during silkworm domestication (Yu *et al*. 2011, Xiang *et al*. 2018). Pigmentation patterning genes are also likely to be associated with the body color differences between *B. mori* and *B. mandarina*. The wild-type *B. mori* larvae are largely white but exhibit melanic spots (*i*.*e*. eye spots, crescent spots and star spots, Figure 1A). *B. mandarina* larvae, on the other hand, are substantially darker, and exhibit extensive dorsal pigment patterning that includes banding and spots (Figure 1A).

In *B. mori*, distinct alleles of the genetic locus, *p*, encode at least 15 different larval markings such as spots, stripes, and banding (Yoda *et al*. 2014). The gene underlying allelic variation at *p, apontic-like* (*apt-like*), encodes a transcription factor that is likely to regulate the expression of melanin synthesis pathway genes such as *yellow-y, ebony, TH, Dopa decarboxylase* (*DDC*) and *laccase 2* (Futahashi *et al*. 2008, Yoda *et al*. 2014) (Figure 1B). While *apt-like* has been implicated in pigmentation differences among *B. mori* strains, it”s potential role in depigmentation of *B. mori* during domestication is not known. A hybrid strain (semiconsomic T02), in which chromosome 2 of *B. mandarina* has been substituted into the genomic background of *B. mori*, exhibits a phenotype similar to that of the *B. mori* allele *moricaud* (*p*^*M*^) (Fujii *et al*. 2021). Since *apt-like* resides on chromosome 2, dorsal pigmentation patterning on the larvae of *B. mandarina* that is absent in *B. mori* was hypothesized to be controlled by the expression of *apt-like* (Yoda *et al*. 2014, Fujii *et al*. 2021), although direct evidence is lacking.

The genetic analysis of domestication-associated loss of pigmentation in *B. mori* has been challenging because deficiencies in candidate genes can result in lethality. For example, in the fruit fly, *Drosophila melanogaster, TH*-deficient (*pale*) mutants die at embryonic stage (Neckameyer and White 1993). In *B. mori, TH*-deficient mutants (*sch lethal, sch*^*l*^) and RNAi-mediated knockdowns of *TH* are both lethal at embryonic stage (Liu *et al*. 2010). In addition, *apt* mutants in *D. melanogaster* die at embryonic stage (Eulenberg and Schuh 1997, Gellon *et al*. 1997), and RNAi-mediated knockdown of *apt-like* in *B. mori* embryos results in death before hatching (Yoda *et al*. 2014).

The issue of lethality is likely to continue to impede the genetic analysis of pigmentation loss and other domestication-associated traits in *B. mori*. Recently, several candidate genes associated with domestication in *B. mori* have been identified using quantitative trait locus (QTL) mapping, including silk production (Li *et al*. 2017, Fang *et al*. 2020), larval climbing ability (Wang, Lin, *et al*. 2020), and mimicry (Wang, Lin, *et al*. 2020). In addition, a recent population genetic analysis identified 300 candidate genes as targets of recent selection in *B. mori*, some of which are likely to be associated with silk production and voltinism (Xiang *et al*. 2018). Despite these efforts, reverse genetic approaches such as targeted gene knockouts and editing (Takasu *et al*. 2010, 2013, Wang *et al*. 2013), gene silencing (Quan *et al*. 2002), or transgenesis (Tamura *et al*. 2000) will likely be required to fully understand the function of these candidate genes and their potential contribution to domestication-associated traits in *B. mori*. Notably, the functions of domestication candidate genes have been tested by reverse genetics approaches in *B. mori* (Xiang *et al*. 2018) but these approaches, to our knowledge, have not yet been implemented in *B. mandarina*.

Here we use CRISPR/Cas9-targeted knockouts of two candidate pigmentation genes in two distinct contexts. First, we demonstrate the feasibility of CRISPR/Cas9-targeted knockouts in *B. mandarina* by generating a homozygous *yellow-y* knockout strain. Next, to circumvent the lethal effects of knocking out a second candidate gene, *apt-like*, we use *allele-specific* CRISPR/Cas9-targeted knockouts in *B. mori* × *B. mandarina* F_1_ hybrids. The latter experiment also comprises a test (the “reciprocal hemizygosity test”, Steinmetz *et al*. 2002, Stern 2014) of the contribution of *apt-like* evolution to the domestication-associated loss of pigmentation in *B. mori*.

## Results and discussion

### A CRISPR/Cas9-targeted knockout of *yellow-y* in *B. mandarina*

To demonstrate the feasibility of CRISPR/Cas9-targeted knockouts in *B. mandarina*, we focused on known pigmentation-related genes. *TH* is among the most compelling candidate genes in the melanin synthesis pathway that seems likely to underlie domestication-associated loss of pigmentation in *B. mori* (Yu *et al*. 2011, Xiang *et al*. 2018). However, *TH* knockout mutants are predicted to be lethal (Neckameyer and White 1993, Liu *et al*. 2010), rendering them difficult to study. Thus, we decided to target *yellow-y* (Figure 1B), a melanin synthesis gene that is downstream of *TH* and functions in melanin synthesis in wide range of insects including *B. mori* and other Lepidoptera and is predicted to be non-essential (Futahashi *et al*. 2008, Zhang *et al*. 2017, Chen *et al*. 2018, Matsuoka and Monteiro 2018, Liu *et al*. 2020, Wang, Huang, *et al*. 2020, Han *et al*. 2021, Shirai *et al*. 2021). After confirming the coding sequence annotation (CDS) of the *B. mandarina yellow-y* gene, we designed a unique CRISPR-RNA (crRNA) target site in exon 2 (Figure 2A, Table S1).

**Figure 2.**
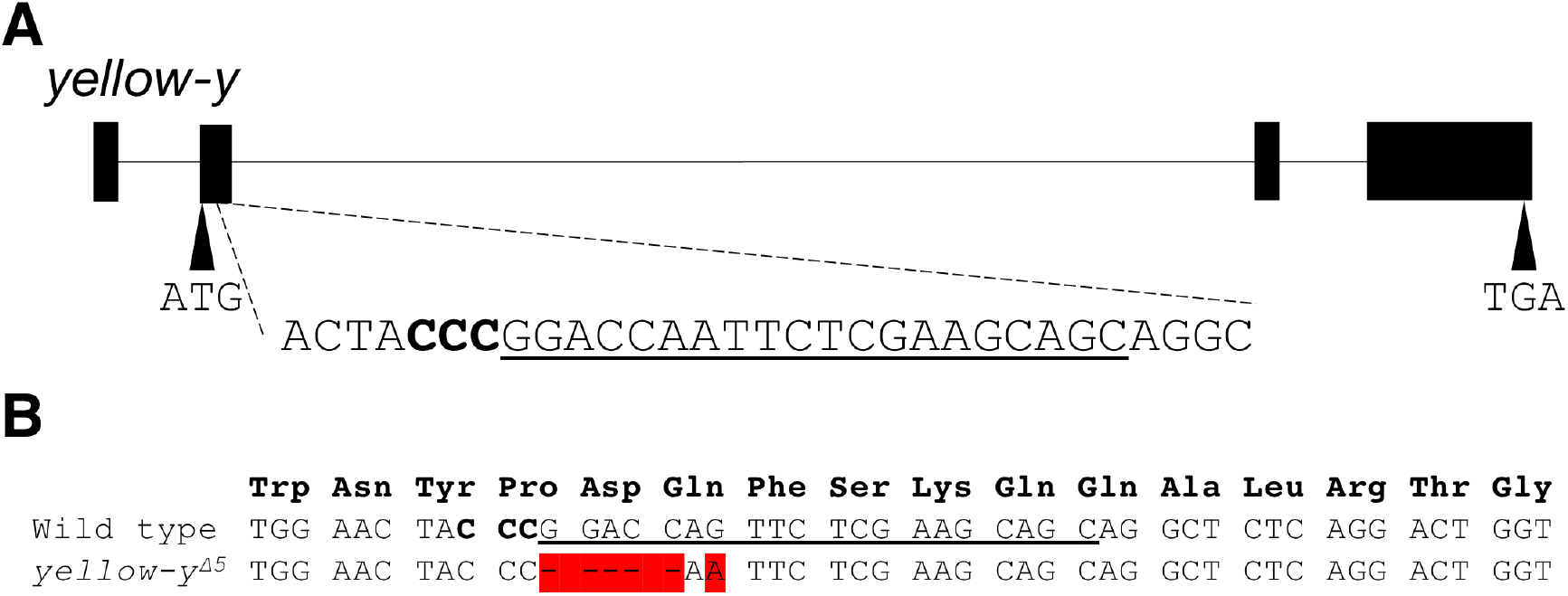
The nature of the lesion in the *yellow-y* knockout (*yellow-y*^Δ15^) in *B. mandarina*. (A)Gene structure of *yellow-y* and the selected crRNA target site. The target sequence is underlined and the protospacer adjacent motif (PAM) site is shown in bold letters. (B) Alignment of the *yellow-y* gene sequences surrounding the crRNA target site from wild type, and G_2_ *B. mandarina* individuals that are homozygous for a 5-base pair deletion and associated single nucleotide mutation. The target sequence is underlined, and the PAM site (CCN) is indicated in bold letters.

In *B. mori*, CRISPR/Cas9-targeted genome editing requires microinjection into non-diapausing eggs (Kanda and Tamura 1994). We obtained non-diapausing eggs by rearing *B. mandarina* larvae under 16 h-light/8 h-dark conditions (Kobayashi 1990). We then injected a mixture of crRNA, trans-activating crRNA (tracrRNA), and Cas9 into 336 *B. mandarina* embryos. Among 16 hatched larvae (G_0_ generation), nine grew to adult moths (Table 1). We crossed six G_0_ adults with wild-type moths and obtained generation 1 (G_1_) eggs. Using a heteroduplex mobility assay on the PCR products from G_1_ embryos, we confirmed that mutations were introduced at the target site in five of the six G_1_ broods (Table 1, Figure S1), showing that CRISPR/Cas9-induced mutations of *yellow-y* were heritable. We then crossed G_1_ siblings with each other and obtained *yellow-y* homozygous knockout individuals carrying a five-nucleotide deletion followed by a single-nucleotide substitution (Figure 2B), which results in a frame-shift and premature stop codon. We designated this mutant allele *yellow-y*^*Δ5*^ and used it for further analyses.

**Table 1.** Efficiency of CRISPR/Cas9-targeted knockout in *B. mandarina* targeting *yellow-y*.

| # eggs injected | # hatched larvae | # adults | # G<sub>1</sub> broods | # G<sub>1</sub> broods carrying mutant alleles |
| --- | --- | --- | --- | --- |
| 336 | 16 | 9 | 6 | 5 |

*B. mandarina yellow-y*^*Δ5*^ homozygotes hatched normally and their development was comparable to that of wild-type individuals (Figure 3). In homozygous *yellow-y*^*Δ5*^ neonate larvae, the larval integument and the head capsule are reddish brown instead of the normal black (Figure 3A) and in final instar larvae, spots and dorsal pigmentation patterns are lighter than that of the wild type (Figure 3B). Later in development, the pupal integument of homozygous *yellow-y*^*Δ5*^ mutants exhibits reddish color instead of the normal black (Figure 3C), and the body and wing spot markings of *yellow-y*^*Δ5*^ adult moths are lighter than that of wild-type (Figure 3D). Further, the phenotypes of heterozygous *+/yellow-y*^*Δ5*^ individuals were comparable to that of wild type (data not shown). Together, these observations suggest that, as observed in *B. mori* (Futahashi *et al*. 2008), *yellow-y* is a non-essential gene contributing to melanin pigment synthesis in *B. mandarina* and loss-of-function is recessive.

**Figure 3.**
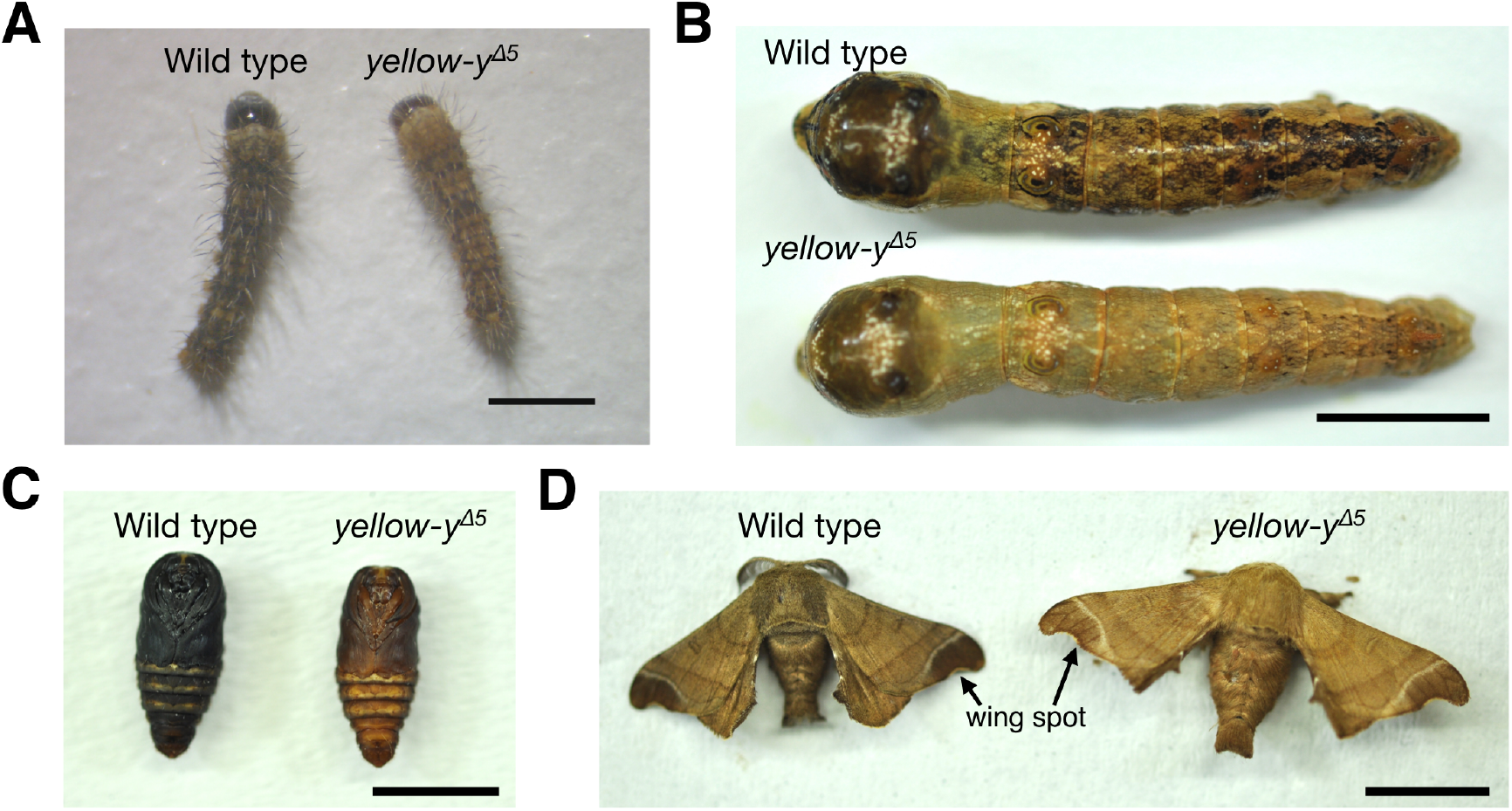
The phenotypes of wild-type and *yellow-y* knockout mutants of *B. mandarina*. Representative (A) neonate larvae, (B) final-instar larvae, (C) male pupae and (D) adult male moths of wild-type (left or top) and *yellow-y*^*Δ5*^ (right or bottom) *B. mandarina*. Arrows indicate wing spot markings in adult moths. Scale bars: 1 mm (A) or 1 cm (B-D).

Comparative data from other species suggests that *yellow-y* functions differently among lepidopteran species. For example, while *yellow-y* loss-of-function is also found to be recessive in several other lepidopteran species (Liu *et al*. 2020, Wang, Huang, *et al*. 2020, Han *et al*. 2021, Shirai *et al*. 2021), it is dominant in the black cutworm, *Agrotis ipsilon* (Chen *et al*. 2018). Further, unlike *Bombyx, yellow-y* knockouts in *Agrotis* are susceptible to dehydration (Chen *et al*. 2018) and mutants in *Spodoptera* exhibit defects in body development, copulation, oviposition, and hatchability (Liu *et al*. 2020, Han *et al*. 2021, Shirai *et al*. 2021). These observations suggest that although the function of *yellow-y* in melanin synthesis is conserved, additional *yellow-y* functions might be diverged among lepidopteran species. *yellow* genes are a rapidly evolving gene family, and loss or duplication of some *yellow* genes have been observed in Lepidoptera (Chen *et al*. 2018, Liu *et al*. 2020, Han *et al*. 2021, Shirai *et al*. 2021). It is possible that functions of *yellow-y* outside of melanin synthesis in *Bombyx* are compensated for by other *yellow* paralogs. In Lepidoptera, the function of most *yellow* genes remained unknown except for *yellow-y* (Futahashi *et al*. 2008, Zhang *et al*. 2017, Chen *et al*. 2018, Matsuoka and Monteiro 2018, Liu *et al*. 2020, Wang, Huang, *et al*. 2020, Han *et al*. 2021, Shirai *et al*. 2021), *yellow-e* (Ito *et al*. 2010), *yellow-d* (Zhang *et al*. 2017) and *yellow-h2/3* (Zhang *et al*. 2017). Further studies of *yellow* genes including *yellow-y* in diverse insect species are required to understand the functions and evolution of *yellow* paralogs.

While our results demonstrate the feasibility of genome editing in *B. mandarina*, these experiments are challenging due to low post-injection hatchability rates (4.8% in our experiment). A recent study showed that the hatchability of *B. mandarina* eggs is generally lower than *B. mori* eggs, even under normal conditions (Zhu *et al*. 2019). To compare the hatchability of *B. mori* and *B. mandarina* embryos after injection, we injected three commonly used buffers (see Methods) and distilled water into embryos of both species and compared hatching rates. We found that the post-injection hatchability of *B. mandarina* embryos (0–8.3 %) is substantially lower than that of *B. mori* (22.9–62.5 %) in all experimental conditions (Table 2), suggesting that *B. mandarina* embryos are more sensitive to injection.

**Table 2.** The hatchability of *B. mori* and *B. mandarina* embryos injected with an injection buffer 1 (Yamaguchi *et al*. 2011), injection buffer 2 (Tamura *et al*. 2000), PBS buffer and distilled water.

| Treatment | Composition | *B. mori* |  | *B. mandarina* |  |
| --- | --- | --- | --- | --- | --- |
|  |  | # injected eggs | # hatched larvae | # injected eggs | # hatched larvae |
| Injection buffer 1 | 100 mM KOAc, 2 mM Mg(OAc)<sub>2</sub>, 30 mM HEPES-KOH; pH 7.4 | 48 | 11 | 48 | 0 |
| Injection buffer 2 | 0.5 mM phosphate buffer (pH 7.0), 5 mM KCl | 48 | 30 | 48 | 4 |
| PBS buffer | 137 mM NaCl, 2.7 mM KCl, 10 mM Na<sub>2</sub>HPO<sub>4</sub>, 1.76mM KH<sub>2</sub>PO<sub>4</sub>; pH 7.4 | 48 | 23 | 48 | 1 |
| Distilled water | — | 48 | 17 | 48 | 2 |

### *Allele-specific* knockouts of *apt-like* in interspecific F_1_ hybrids

Knockouts of some pigmentation pathway genes, such as *apt-like* and *TH*, are predicted to be lethal (Neckameyer and White 1993, Eulenberg and Schuh 1997, Gellon *et al*. 1997, Liu *et al*. 2010, Yoda *et al*. 2014). Considering this obstacle to the study of essential genes, together with the observation (above) of reduced hatchability of injected *B. mandarina* embryos, we instead opted for the alternative strategy of injecting *B. mori* × *B. mandarina* F_1_ hybrids. Specifically, we conducted an *allele-specific* CRISPR/Cas9-targeted knockout of *apt-like* in F_1_ embryos (from crosses between *B. mori* females and *B. mandarina* males). We reasoned that, because the components of these F_1_ eggs is derived from the *B. mori* mother, these embryos should have post-injection hatchability similar to that of *B. mori*. We designed species-specific crRNA targeting *apt-like*, for which the targeted protospacer adjacent motif (PAM) sequence is only present in either *B. mori-* or *B. mandarina*-derived sequence (Methods, Figure 4, Table S1). For each targeted allele, we injected a mixture of crRNA, tracrRNA and Cas9 into 48 F_1_ embryos. To confirm that mutations were specifically introduced into the targeted allele, we extracted genomic DNA from adult legs and PCR-amplified the target sites. We then carried out heteroduplex mobility assays using a microchip electrophoresis system (Ota *et al*. 2013, Ansai *et al*. 2014), which show that various mutations were introduced into the *apt-like* target sequence in both of the *allele-specific* knockout series (Figure S2). The PCR products obtained from two representative individuals of each *allele-specific* knockout series were then cloned and sequenced (Figure 4). The sequences of cloned PCR products confirmed that various mutations were introduced into the target sites in both knockout series, some of which cause frameshifts and associated premature stop codons. All mutations detected in this experiment were specifically introduced only into the targeted allele.

**Figure 4.**
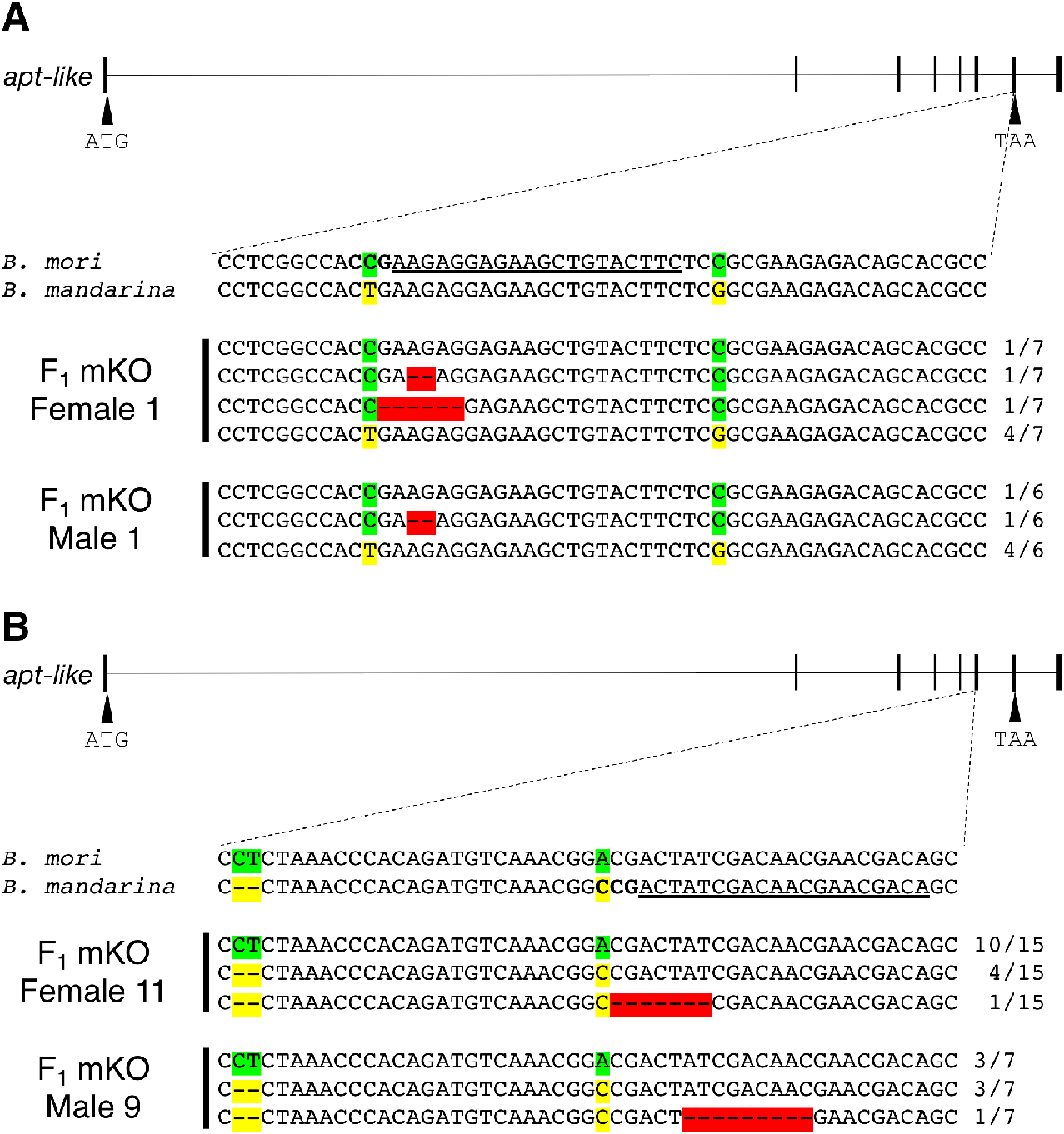
Mutations introduced by *allele-specific apt-like* mosaic knockout (mKO) in F_1_ hybrids. Annotation and inset alignments of the *apt-like* gene in both species with selected crRNA target sites whose PAM sequences are only present in (A) *B. mori* or *(A) B. mandarina*. Two *allele-specific* knockout individuals were sequenced for each target. The target sequences are underlined, and the PAM sequences are shown in bold letters. *B. mori* specific SNPs and *B. mandarina* specific single nucleotide variants are highlighted with green and yellow shadings, respectively. Mutations introduced by CRISPR/Cas9 system are highlighted with red shading. Numbers on the right edge indicate the numbers of the clones identified among all cloned and sequenced PCR products.

In addition to spots, normal F_1_ larvae have a dark body with darker greyish brown banding covering wide range of the dorsal surface, similar to the pattern for *B. mandarina* (Figure 1A, Figure S3). As predicted, post-injection hatchability of F_1_ embryos was high whether targeting the *B. mori* (60%) or the *B. mandarina* (66%) allele. In the *apt-like* knockout series targeting the *B. mori* allele, all of the 26 larvae that survived to the fifth instar stage exhibited normal body color (Table 3, Figure 5). In contrast, for the *apt-like* knockout series targeting the *B. mandarina* allele, 24 of the 29 larvae that survived to the fifth instar stage exhibited white patches on dorsal pigmentation pattern banding that varied in size (Table 3, Figure 5, Figure S4). This result establishes a role for *apt-like* in body pigment formation and that the *B. mandarina*-derived *apt-like* allele is dominant with respect to this trait. Notably, however, the pigmentation of F_1_ pupae and adult stages of the knockout series targeting the *B. mori* or the *B. mandarina* allele both exhibited normal body color (data not shown). Additionally, larval spots, which are also predicted to be controlled by *apt-like* (Yoda *et al*. 2014), were also not affected the F_1_ series targeting either the *B. mori* or the *B. mandarina* allele (Figure 5, Figure S4). Since both wild-type (+^*p*^) *B. mori* and *B. mandarina* larvae exhibit spots (Figure 1A), we conclude that the *B. mori* or the *B. mandarina*-derived *apt-like* alleles are both sufficient to direct the formation of larval spots.

**Table 3.** Efficiency of *allele-specific* knockouts of *apt-like* in F_1_ hybrids.

| Targeted allele | # eggs injected | # hatched larvae | # 5th instar larvae | # mosaic 5th instar larvae | # adults (male/female) |
| --- | --- | --- | --- | --- | --- |
| *B. mori* | 48 | 29 | 26 | 0 | 26 (16/10) |
| *B. mandarina* | 48 | 32 | 29 | 24 | 29 (18/11) |

**Figure 5.**
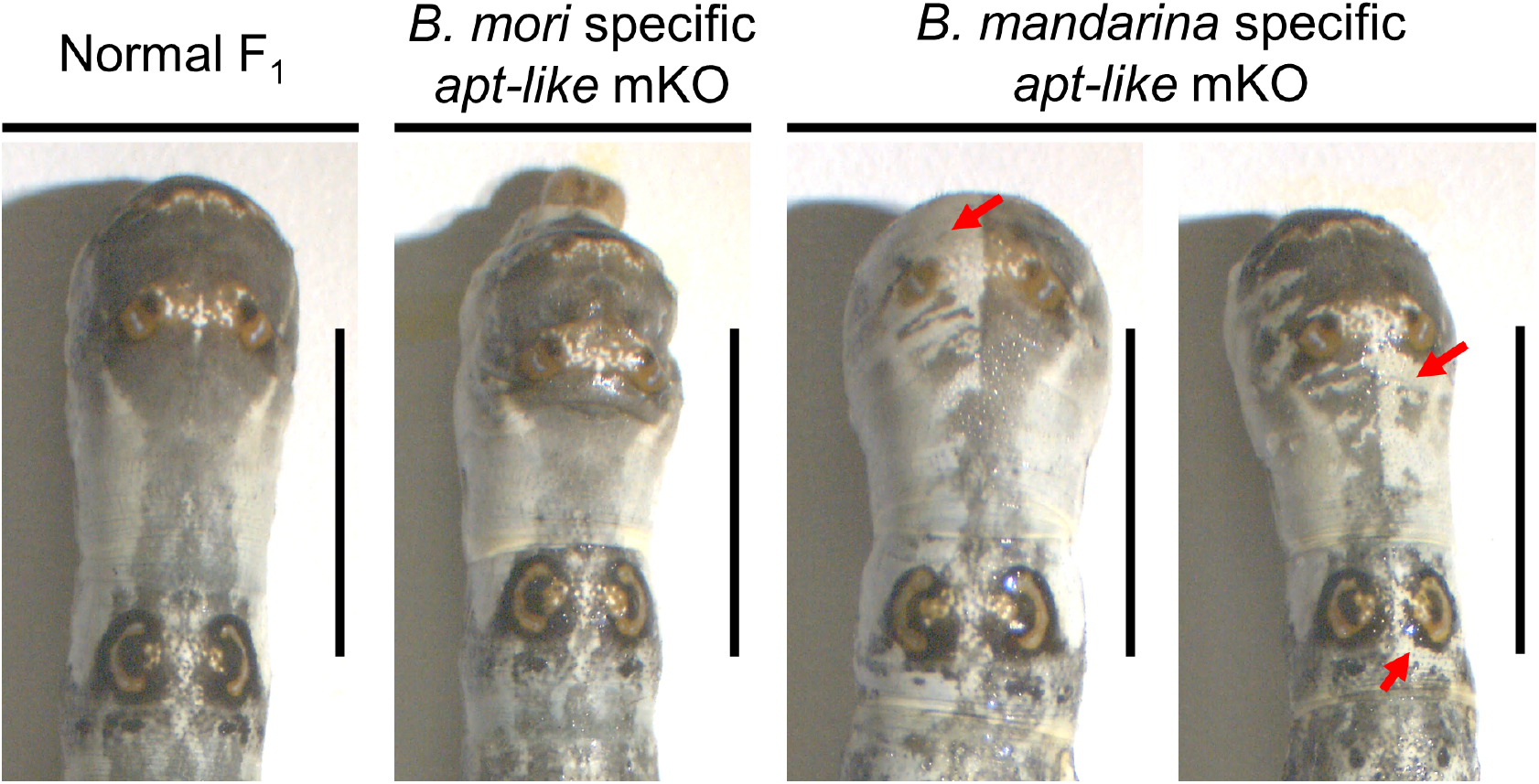
Representative fifth instar larvae of normal F_1_ (left), *allele-specific apt-like* mKOs targeting the *B. mori* allele (middle) and the *B. mandarina* allele (right). Red arrows indicate ectopic white (depigmented) regions. Scale bars: 1 cm.

Our results have implications beyond merely confirming a role for *apt-like* in *B. mandarina* larval pigmentation. The *allele-specific* knockouts of *apt-like* in F_1_ hybrids allows us to compare the phenotype of genetically identical hybrids that differ only at the target locus (a framework called the “reciprocal hemizygosity test”, Steinmetz *et al*. 2002, Stern 2014). As such, we can attribute the loss of pigmentation in the series targeting the *B. mandarina* allele to evolution at the *apt-like* gene in *B. mori*, rather than exclusively at a *trans*-acting factor. The Apt-like proteins of *B. mori* (p50T) and *B. mandarina* (Sakado) differ by only one amino acid substitution: Alanine to Valine at residue 188 (see DDBJ accession numbers LC706749 and LC706750). However, this substitution is not observed in *B. mandarina* collected at different locations (see NCBI accession numbers SRR6111377, SRR6111379, SRR6111381 and SRR6111382), suggesting this is not a fixed amino acid difference between species. This implies that evolution of an *apt-like cis*-regulatory element contributes to the observed phenotypic difference between species.

## Conclusion

Here we demonstrate the utility of CRISPR/cas9 genome editing in *B. mandarina* and *B. mori* × *B. mandarina* F_1_ hybrids to the study the function and evolution of domestication-associated candidate genes. Focusing on two pigmentation-related genes, we show that *apt-like* plays a role in larval body pigmentation patterning (Figure 5), whereas *yellow-y* plays a more general role in pigmentation that is not pattern dependent or specific to developmental life-stage (Figure 3B). These results are consistent with the proposed roles of Apt-like as a transcription factor responsible for larval color patterning, and Yellow-y as an enzyme in the melanin synthesis pathway under control of Apt-like (Futahashi *et al*. 2008, Yoda *et al*. 2014). Further, using the framework of the reciprocal hemizygosity test, we show that *apt-like* has evolved in *B. mori* in a way that has specifically reduced larval body pigmentation, without affecting the formation of larval spots or adult body pigmentation.

Despite being a powerful tool to study gene function and evolution, the *allele-specific* knockout approach has several limitations. First, it requires sequence variants distinguishing the parents of the F_1_ hybrid that result in a PAM-site that is only present in one of two species, limiting the potential to design *allele-specific* targets. However, this limitation may be overcome by using Cas proteins that recognize different PAM-sites (Leenay and Beisel 2017). For example, while the *Streptococcus pyogenes* Cas9 (*Sp*Cas9) protein has been used here, the Cas12a, which recognizes a distinct PAM sequence, has also been implemented in *B. mori* (Dong *et al*. 2020). In addition, recent studies have reported that *Sp*Cas9 can be modified to recognize alternative PAM sequences (Kleinstiver *et al*. 2015, 2016). A second limitation is that, given the mosacism of knockouts in G_0_ individuals, one cannot exclude the possibility of false-negative phenotyping results. The cleavage efficiency of the CRISPR/Cas9 system is affected by several features, such as the sequences of PAM-distal and PAM-proximal regions of the guide RNA, the genomic context of the targeted DNA, as well as GC-content and secondary structure of the guide RNA (Liu *et al*. 2016). To minimize this problem, one can screen a large number of G_0_ individuals and confirm that mutations were introduced with high efficiency (as in Figure 4 and Figure S2).

Despite these limitations, our results highlight several advantages of *allele-specific* knockouts in the F_1_ over knockouts in *B. mandarina*. First, our results show that *B. mori* (female) × *B. mandarina* (male) F_1_ embryos are substantially more tolerant to injection compared to *B. mandarina*. Second, *allele-specific* knockouts in the F_1_ permit the study of essential genes (such as *apt-like*) at which knockouts are expected to be homozygous lethal and recessive with respect to the phenotype. Finally, *allele-specific* CRISPR/Cas9-targeted knockouts in F_1_ hybrids have the added utility of identifying loci that have diverged in function between *B. mori* and *B. mandarina*, and contributing to domestication-related traits in *B. mori* using the framework of the reciprocal hemizygosity test. Thus, our study showcases the multifaceted utility of *allele-specific* knockouts in F_1_ hybrids in the study of gene function and evolution.

## Experimental procedures

### Insects

The *B. mori* strain p50T, a single-paired descendant of individuals of strain p50 (a derivative from Daizo; https://shigen.nig.ac.jp/silkwormbase/ViewStrainDetail.do?name=p50), is maintained at our laboratory. The *B. mandarina* strain, Sakado, was originally collected in Sakado-city, Saitama, Japan, in 1982. Since then, it has been maintained at our laboratory by sib-mating. All larvae were reared on fresh mulberry leaves or artificial diet (SilkMate PS, NOSAN) under continuous 12 h-light/12 h-dark conditions at 25 °C with the exceptions described below. For injections in to *B. mandarina* embryos, we obtained non-diapausing eggs by rearing *B. mandarina* larvae under continuous long-day conditions (16 h-light/8 h-dark) at 25 °C (Kobayashi 1990). We then collected eggs in crosses between emerged adults. To generate *B. mori* × *B. mandarina* F_1_ hybrid embryos for injection, we first incubated *B. mori* eggs at 15 ºC under continuous darkness (Kogure 1933). The hatched larvae were then reared under continuous 16 h-light/8 h-dark condition at 25 °C and females were crossed to *B. mandarina* males.

### Confirmation of the *yellow-y* and *apt-like* coding sequences in *B. mandarina*

Total RNA was extracted from the integument of fourth instar *B. mandarina* larvae using TRIzol (Thermo Fisher Scientific). Complementary DNA (cDNA) was reverse transcribed from total RNA using TaKaRa RNA PCR Kit (TaKaRa). Reverse transcriptase-PCR was performed using KOD One polymerase (TOYOBO). PCR products were cloned into pGEM-T Easy Vector (Promega) and Sanger-sequenced using the FASMAC sequencing service (Kanagawa, Japan).

### Knockout of *yellow-y* in *B. mandarina*

An unique crRNA target sequence in the *B. mori* genome was selected using CRISPRdirect (https://crispr.dbcls.jp) (Table S1) (Naito *et al*. 2015). The uniqueness of the target sequence in the *B. mandarina* genome was then confirmed by performing blastn at SilkBase (http://silkbase.ab.a.u-tokyo.ac.jp). A mixture of crRNA, tracrRNA and Cas9 Nuclease protein NLS (600 ng/µL; NIPPON GENE) in injection buffer (100 mM KOAc, 2 mM Mg(OAc)_2_, 30 mM HEPES-KOH; pH 7.4) was injected into each embryo within 3 h after oviposition (Yamaguchi *et al*. 2011).

The injected embryos (G_0_ generation) were incubated at 25 °C in a humidified Petri dish until hatching. Adult G_0_ moths were crossed with wild-type *B. mandarina*, and G_1_ eggs were obtained. To detect heritable CRISPR/Cas9-induced mutations, ten G_1_ eggs were collected into one tube, and genomic DNA was prepared using the HotSHOT method (Truett *et al*. 2000). The region containing the target site of *yellow-y* crRNA was PCR-amplified using KOD One polymerase (TOYOBO). Mutations at the target site were detected by heteroduplex mobility assay using the MultiNA microchip electrophoresis system (SHIMAZU) with the DNA-500 reagent kit (Ota *et al*. 2013, Ansai *et al*. 2014).

Adult G_1_ moths from broods with heritable CRISPR/Cas9-induced mutations were crossed with each other to obtain homozygous knockout mutants (G_2_). To confirm CRISPR/Cas9-induced mutations, we prepared genomic DNA, PCR-amplified the target region and detected mutations as described above using G_2_ adult moths. To determine the precise nature of insertions, deletions and substitutions, PCR products obtained from G_2_ individuals were directly Sanger-sequenced using the FASMAC sequencing service (Kanagawa, Japan).

### Comparison of post-injection hatchability

Three commonly used buffers (injection buffer 1 (Yamaguchi *et al*. 2011), injection buffer 2 (Tamura *et al*. 2000), PBS buffer) and distilled water into embryos of *B. mori* or *B. mandarina* within 3 h after oviposition. The injected embryos were incubated at 25 °C in a humidified Petri dish until hatching.

### *Allele-specific* gene knockouts in F_1_ hybrids

A PAM sequence is necessary for target recognition and following DNA cleavage in CRISPR/Cas system (Hsu *et al*. 2013, Anders *et al*. 2014), and *Sp*Cas9, which we used in this study, recognizes 5”-NGG-3” as PAM. We specifically targeted *Sp*Cas9 PAM-sites that differed in sequence between *B. mori* and *B. mandarina* (Figure 4, Table S1), allowing *allele-specific* DNA cleavage and gene knockout (Courtney *et al*. 2015, Christie *et al*. 2017). A mixture of crRNA, tracrRNA and Cas9 was injected to F_1_ embryos as described above. To evaluate CRISPR/Cas9 cleavage efficiency, we extracted genomic DNA from G_0_ adult legs and PCR-amplified the target region as described above. PCR products were cloned into pGEM-T Easy Vector and Sanger-sequenced using an ABI3130xl genetic analyzer (Applied Biosystems).

## Acknowledgements

We are grateful to Katsuya Satta and Hisashi Tobita for assistance with the experiments. We are also grateful to Professor Susumu Katsuma for helpful discussions. We thank the Institute for Sustainable Agro-ecosystem Services, The University of Tokyo, for facilitating the mulberry cultivation and the Biotron Facility at the University of Tokyo for rearing the silkworms. This work was supported by JSPS KAKENHI grant number JP20J22954 to KT and JP20H02997 to TK.

## Data Availability

Full-length coding sequences of *B. mori yellow-y, B. mandarina yellow-y, B. mori apt-like* and *B. mandarina apt-like* are available on the DDBJ under accession numbers of LC706747, LC706748, LC706749 and LC706750, respectively.

## Author contributions

KT, PA and TK designed the study. KT performed most of the experiments. KT wrote the manuscript with intellectual input from TK and PA. All authors edited and approved the final version of the manuscript and agree to be accountable for all aspects of the work.

---

**Table S1.**
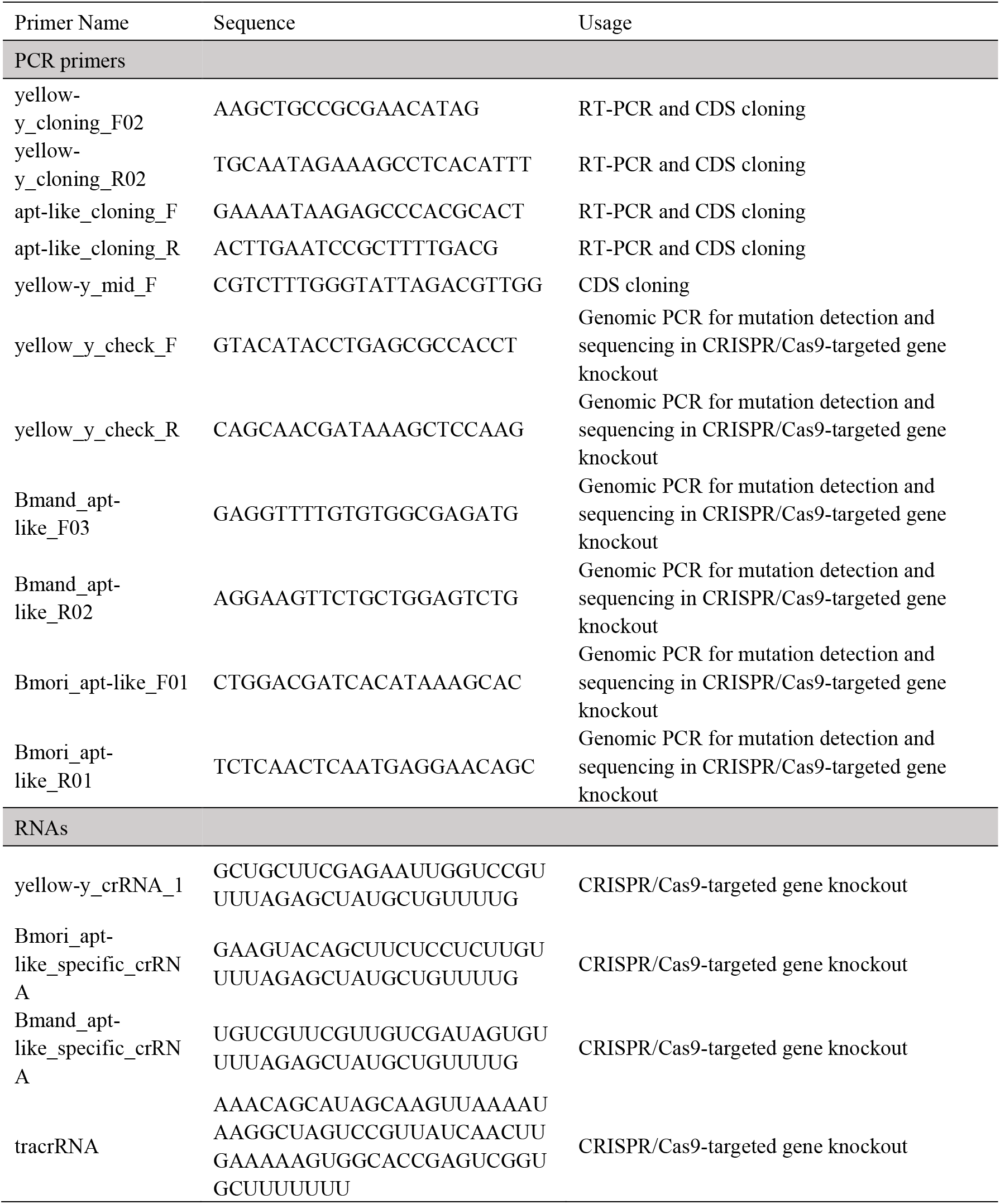
PCR primers and RNA sequences used in this study.

**Figure S1.**
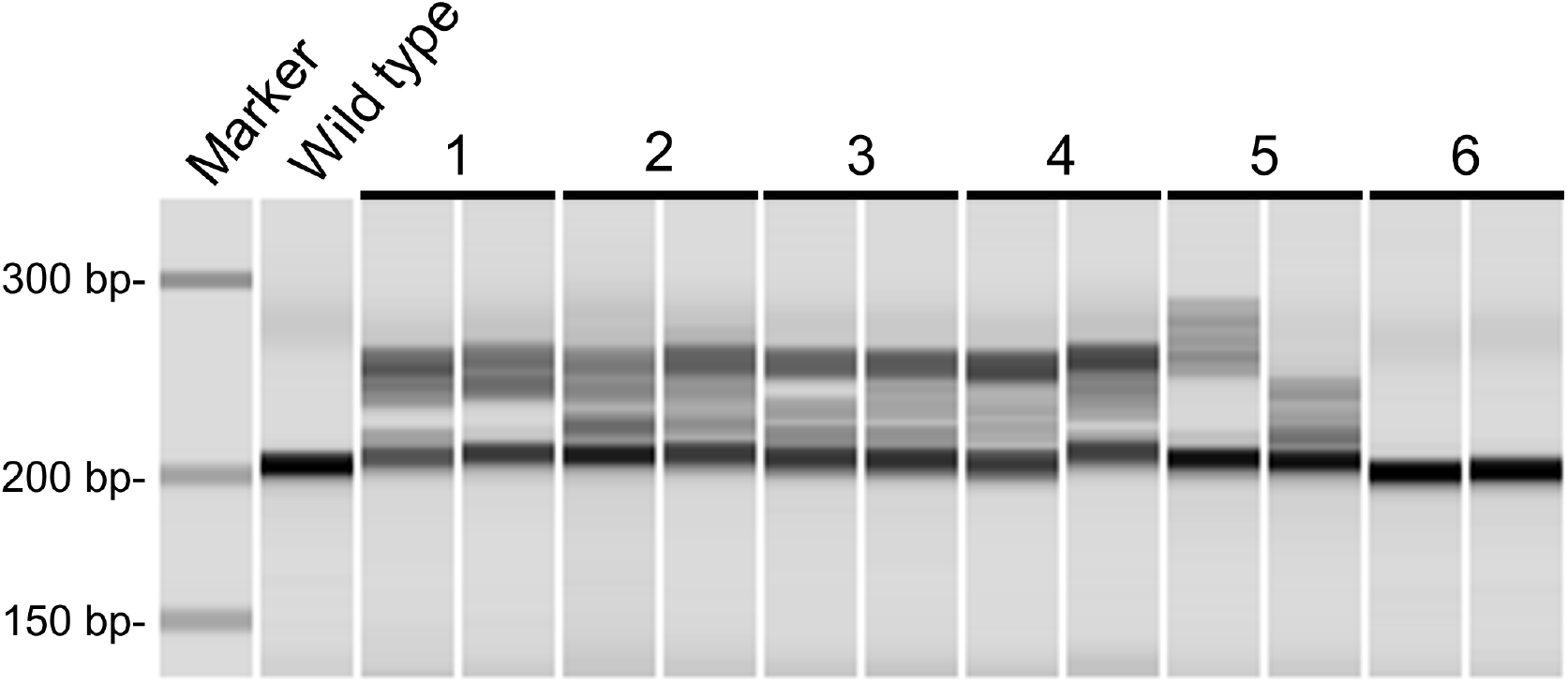
Detection of mutations at the *yellow-y* crRNA target site using a heteroduplex mobility shift assay. Two intervals were prepared for each G_1_ brood. Ten eggs were collected into one tube, and genomic DNA was prepared. The region containing the target site of *yellow-y* crRNA was PCR-amplified. Multiple heteroduplex bands caused by insertion/deletion mismatches were observed indicating the presence of DNA lesions (insertions/deletions and/or associated nucleotide variants).

**Figure S2.**
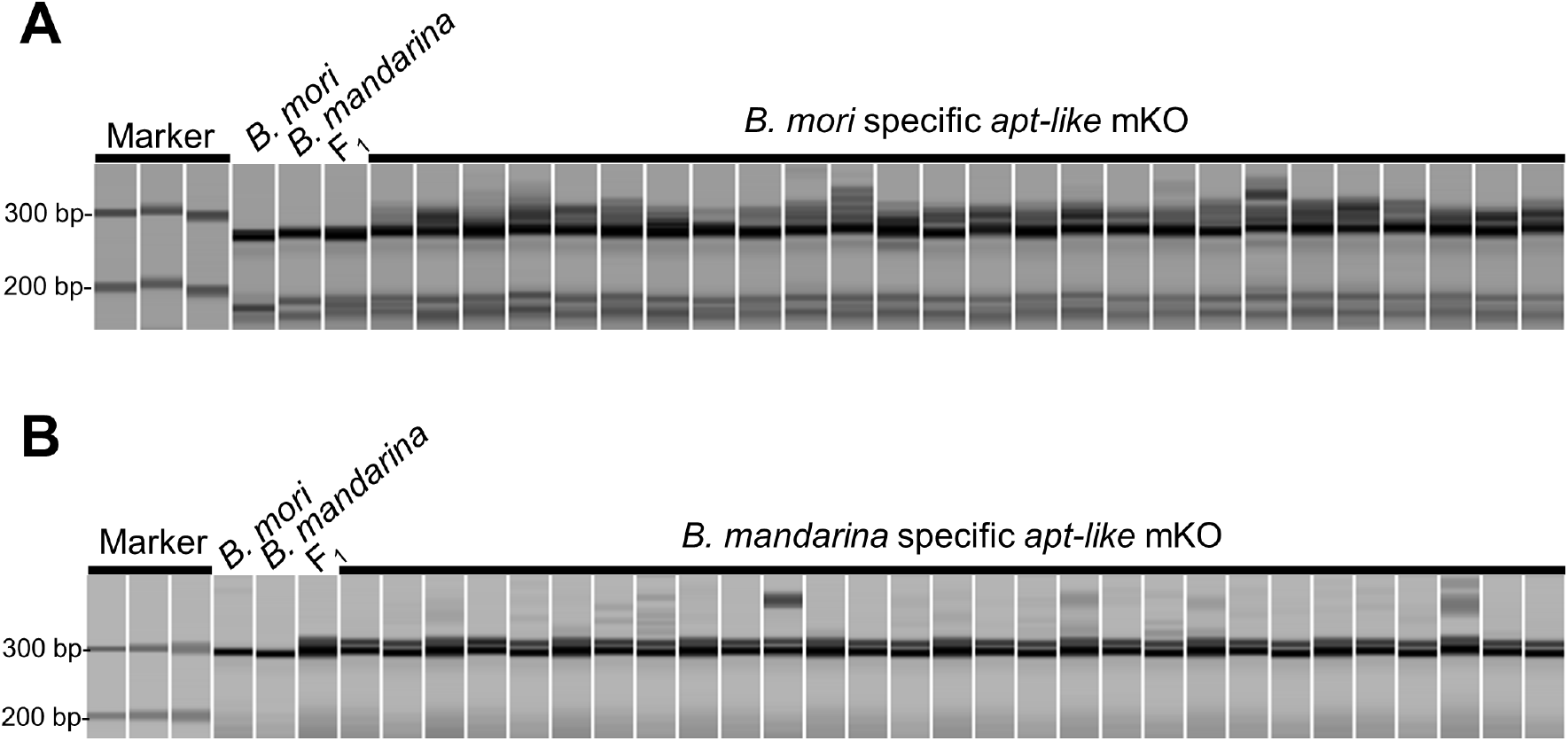
Detection of mutations at *apt-like* crRNA target sites using a heteroduplex mobility shift assay. The region containing the target site of (A) *B. mori* specific *apt-like* crRNA and (B) *B. mandarina* specific *apt-like* crRNA was PCR-amplified using DNA prepared from G_0_ adults” legs. Multiple heteroduplex bands caused by insertion/deletion mismatches and associated nucleotide variants were observed in G_0_ mosaics.

**Figure S3.**
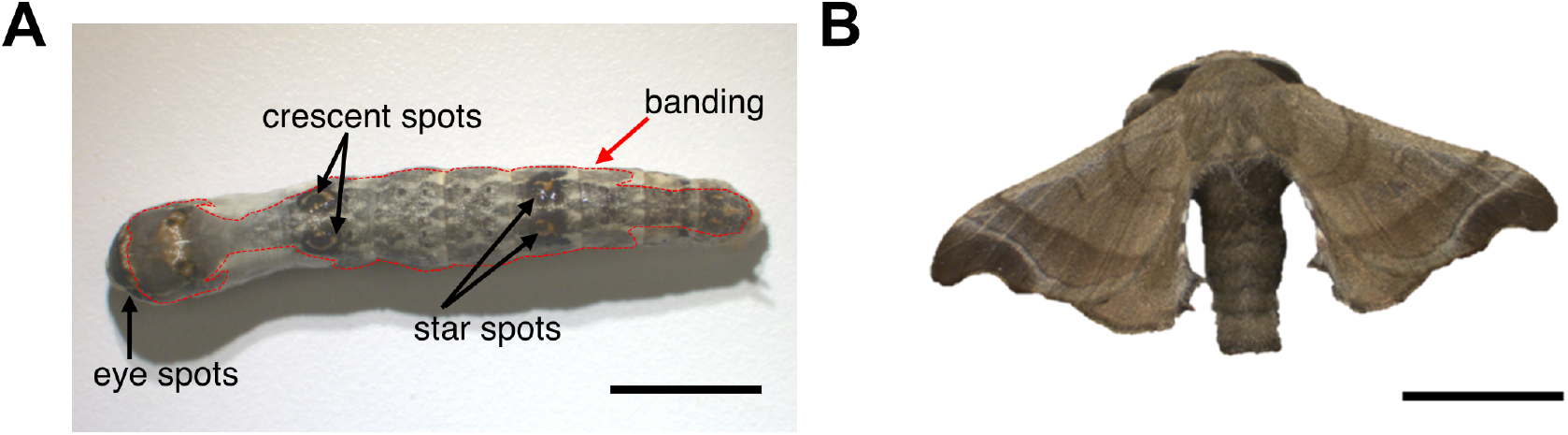
A representative (A) fifth instar larva and (B) adult male moth of normal F_1_ (*B mori* female × *B. mandarina* male). Black arrows indicate larval spot markings and red dotted lines outline the banding. Normal F_1_ larvae have a dark body with darker greyish brown banding covering wide range of the dorsal surface similar to *B. mandarina*. The names of spots were referred to Yoda *et al*. (2014). Scale bars: 1 cm.

**Figure S4.**
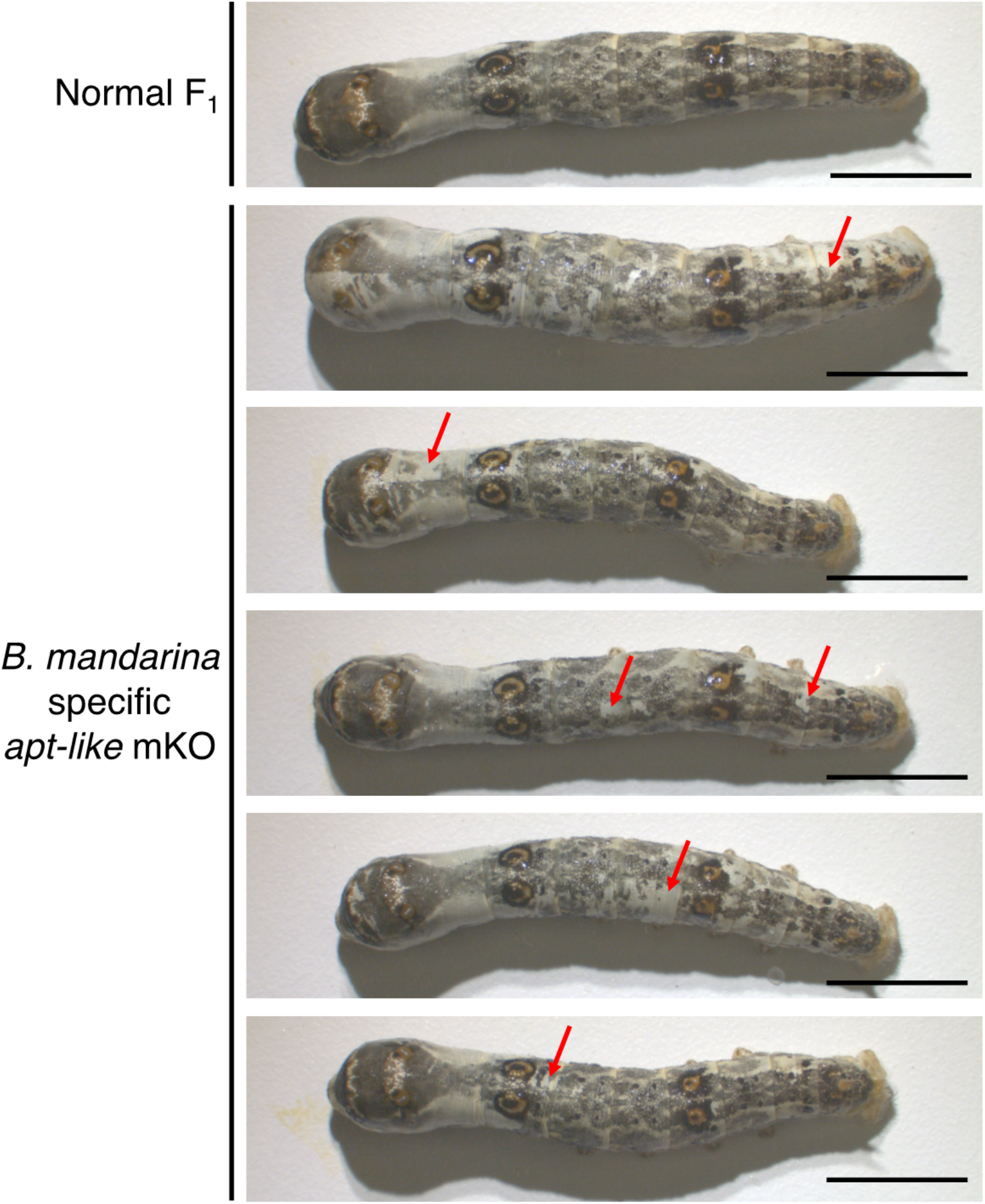
Fifth instar larvae of normal F_1_ and *B. mandarina* specific *apt-like* mKO. Red arrows indicate ectopic white (depigmented) regions. Crescent and star spots were not depigmented in all larvae. Scale bars: 1 cm.

## References

Anders, C., Niewoehner, O., Duerst, A., and Jinek, M., 2014. Structural basis of PAM-dependent target DNA recognition by the Cas9 endonuclease. Nature, 513 (7519), 569–573.

Ansai, S., Inohaya, K., Yoshiura, Y., Schartl, M., Uemura, N., Takahashi, R., and Kinoshita, M., 2014. Design, evaluation, and screening methods for efficient targeted mutagenesis with transcription activator-like effector nucleases in medaka. Development, Growth & Differentiation, 56 (1), 98–107.

Chen, X., Cao, Y., Zhan, S., Zhang, Y., Tan, A., and Huang, Y., 2018. Identification of yellow gene family in Agrotis ipsilon and functional analysis of Aiyellow-y by CRISPR/Cas9. Insect Biochemistry and Molecular Biology, 94, 1–9.

Christie, K.A., Courtney, D.G., DeDionisio, L.A., Shern, C.C., De Majumdar, S., Mairs, L.C., Nesbit, M.A., and Moore, C.B.T., 2017. Towards personalised allele-specific CRISPR gene editing to treat autosomal dominant disorders. Scientific Reports, 7 (1), 16174.

Courtney, D.G., Moore, J.E., Atkinson, S.D., Maurizi, E., Allen, E.H.A., Pedrioli, D.M.L., McLean, W.H.I., Nesbit, M.A., and Moore, C.B.T., 2015. CRISPR/Cas9 DNA cleavage at SNP-derived PAM enables both in vitro and in vivo KRT12 mutation-specific targeting. Gene Therapy, 23 (1), 108–112.

Dong, Z., Qin, Q., Hu, Z., Zhang, X., Miao, J., Huang, L., Chen, P., Lu, C., and Pan, M., 2020. CRISPR/Cas12a mediated genome editing enhances Bombyx mori resistance to BmNPV. Frontiers in Bioengineering and Biotechnology, 8, 841.

Eulenberg, K.G. and Schuh, R., 1997. The tracheae defective gene encodes a bZIP protein that controls tracheal cell movement during Drosophila embryogenesis. The EMBO Journal, 16 (23), 7156–7165.

Fang, S., Zhou, Q., Yu, Q., and Zhang, Z., 2020. Genetic and genomic analysis for cocoon yield traits in silkworm. Scientific Reports, 10 (1), 5682.

Fujii, T., Kiuchi, T., Daimon, T., Ito, K., Katsuma, S., Shimada, T., Yamamoto, K., and Banno, Y., 2021. Development of interspecific semiconsomic strains between the domesticated silkworm, Bombyx mori and the wild silkworm, B. mandarina. Journal of Insect Biotechnology and Sericology, 90 (2), 33–40.

Futahashi, R., Sato, J., Meng, Y., Okamoto, S., Daimon, T., Yamamoto, K., Suetsugu, Y., Narukawa, J., Takahashi, H., Banno, Y., Katsuma, S., Shimada, T., Mita, K., and Fujiwara, H., 2008. yellow and ebony are the responsible genes for the larval color mutants of the silkworm Bombyx mori. Genetics, 180 (4), 1995–2005.

Gellon, G., Harding, K.W., McGinnis, N., Martin, M.M., and McGinnis, W., 1997. A genetic screen for modifiers of deformed homeotic function identifies novel genes required for head development. Development, 124 (17), 3321–3331.

Han, W., Tang, F., Zhong, Y., Zhang, J., and Liu, Z., 2021. Identification of yellow gene family and functional analysis of Spodoptera frugiperda yellow-y by CRISPR/Cas9. Pesticide Biochemistry and Physiology, 178, 104937.

Hsu, P.D., Scott, D.A., Weinstein, J.A., Ran, F.A., Konermann, S., Agarwala, V., Li, Y., Fine, E.J., Wu, X., Shalem, O., Cradick, T.J., Marraffini, L.A., Bao, G., and Zhang, F., 2013. DNA targeting specificity of RNA-guided Cas9 nucleases. Nature Biotechnology, 31 (9), 827–832.

Ito, K., Katsuma, S., Yamamoto, K., Kadono-Okuda, K., Mita, K., and Shimada, T., 2010. Yellow-e determines the color pattern of larval head and tail spots of the silkworm Bombyx mori. Journal of Biological Chemistry, 285 (8), 5624–5629.

Kanda, T. and Tamura, T., 1994. Microinjection method for DNA in early embryos of the silkworm, Bombyx mori, using air-pressure. Bulletin of the National Institute of Sericultural and Entomological Science, 2, 31-46 (in Japanese with English summary).

Kleinstiver, B.P., Pattanayak, V., Prew, M.S., Tsai, S.Q., Nguyen, N.T., Zheng, Z., and Joung, J.K., 2016. High-fidelity CRISPR–Cas9 nucleases with no detectable genome-wide off-target effects. Nature, 529 (7587), 490–495.

Kleinstiver, B.P., Prew, M.S., Tsai, S.Q., Topkar, V. V., Nguyen, N.T., Zheng, Z., Gonzales, A.P.W., Li, Z., Peterson, R.T., Yeh, J.-R.J., Aryee, M.J., and Joung, J.K., 2015. Engineered CRISPR-Cas9 nucleases with altered PAM specificities. Nature, 523 (7561), 481–485.

Kobayashi, J., 1990. Effects of photoperiod on the induction of egg diapause of tropical races of the domestic silkworm, Bombyx mori, and the wild silkworm, B. mandarina. Japan Agricultural Research Quarterly, 23 (3), 202–205.

Kogure, M., 1933. The influence of light and temperature on certain characters of the silkworm, Bombyx mori. Journal of the Faculty of Agriculture, Kyushu University, 4 (1), 1–93.

Leenay, R.T. and Beisel, C.L., 2017. Deciphering, communicating, and engineering the CRISPR PAM. Journal of Molecular Biology, 429 (2), 177–191.

Li, C., Tong, X., Zuo, W., Luan, Y., Gao, R., Han, M., Xiong, G., Gai, T., Hu, H., Dai, F., and Lu, C., 2017. QTL analysis of cocoon shell weight identifies BmRPL18 associated with silk protein synthesis in silkworm by pooling sequencing. Scientific Reports, 7 (1), 17985.

Liu, C., Yamamoto, K., Cheng, T.C., Kadono-Okuda, K., Narukawa, J., Liu, S.P., Han, Y., Futahashi, R., Kidokoro, K., Noda, H., Kobayashi, I., Tamura, T., Ohnuma, A., Banno, Y., Dai, F.Y., Xiang, Z.H., Goldsmith, M.R., Mita, K., and Xia, Q.Y., 2010. Repression of tyrosine hydroxylase is responsible for the sex-linked chocolate mutation of the silkworm, Bombyx mori. Proceedings of the National Academy of Sciences of the United States of America, 107 (29), 12980–12985.

Liu, X., Han, W., Ze, L., Peng, Y., Yang, Y., Zhang, J., Yan, Q., and Dong, S., 2020. Clustered regularly interspaced short palindromic repeats/CRISPR-associated protein 9 mediated knockout reveals functions of the yellow-y gene in Spodoptera litura. Frontiers in Physiology, 11, 1643.

Liu, X., Homma, A., Sayadi, J., Yang, S., Ohashi, J., and Takumi, T., 2016. Sequence features associated with the cleavage efficiency of CRISPR/Cas9 system. Scientific Reports, 6 (1), 19675.

Lu, K., Liang, S., Han, M., Wu, C., Song, J., Li, C., Wu, S., He, S., Ren, J., Hu, H., Shen, J., Tong, X., and Dai, F., 2020. Flight muscle and wing mechanical properties are involved in flightlessness of the domestic silkmoth, Bombyx mori. Insects, 11 (4), 220.

Matsuoka, Y. and Monteiro, A., 2018. Melanin pathway genes regulate color and morphology of butterfly wing scales. Cell reports, 24 (1), 56–65.

Naito, Y., Hino, K., Bono, H., and Ui-Tei, K., 2015. CRISPRdirect: software for designing CRISPR/Cas guide RNA with reduced off-target sites. Bioinformatics, 31 (7), 1120–1123.

Neckameyer, W.S. and White, K., 1993. Drosophila Tyrosine hydroxylase is encoded by the pale locus. Journal of Neurogenetics, 8 (4), 189–199.

Ômura, S., 1950. Researches on the behavior and ecological characteristics of the wild silkworm Bombyx mandarina. Bulletin of the Imperial Sericultural Experiment Station, 13 (3), 79-130 (in japanese with English summary).

Ota, S., Hisano, Y., Muraki, M., Hoshijima, K., Dahlem, T.J., Grunwald, D.J., Okada, Y., and Kawahara, A., 2013. Efficient identification of TALEN-mediated genome modifications using heteroduplex mobility assays. Genes to Cells, 18 (6), 450–458.

Quan, G.-X., Kim, I., Kômoto, N., Sezutsu, H., Ote, M., Shimada, T., Kanda, T., Mita, K., Kobayashi, M., and Tamura, T., 2002. Characterization of the kynurenine 3-monooxygenase gene corresponding to the white egg 1 mutant in the silkworm Bombyx mori. Molecular Genetics and Genomics, 267 (1), 1–9.

Shirai, Y., Ohde, T., and Daimon, T., 2021. Functional conservation and diversification of yellow-y in lepidopteran insects. Insect Biochemistry and Molecular Biology, 128, 103515.

Steinmetz, L.M., Sinha, H., Richards, D.R., Spiegelman, J.I., Oefner, P.J., McCusker, J.H., and Davis, R.W., 2002. Dissecting the architecture of a quantitative trait locus in yeast. Nature, 416 (6878), 326–330.

Stern, D.L., 2014. Identification of loci that cause phenotypic variation in diverse species with the reciprocal hemizygosity test. Trends in Genetics, 30 (12), 547–554.

Takasu, Y., Kobayashi, I., Beumer, K., Uchino, K., Sezutsu, H., Sajwan, S., Carroll, D., Tamura, T., and Zurovec, M., 2010. Targeted mutagenesis in the silkworm Bombyx mori using zinc finger nuclease mRNA injection. Insect Biochemistry and Molecular Biology, 40 (10), 759–765.

Takasu, Y., Sajwan, S., Daimon, T., Osanai-Futahashi, M., Uchino, K., Sezutsu, H., Tamura, T., and Zurovec, M., 2013. Efficient TALEN construction for Bombyx mori gene targeting. PLoS ONE, 8 (9), e73458.

Tamura, T., Thibert, C., Royer, C., Kanda, T., Eappen, A., Kamba, M., Kômoto, N., Thomas, J.-L., Mauchamp, B., Chavancy, G., Shirk, P., Fraser, M., Prudhomme, J.-C., and Couble, P., 2000. Germline transformation of the silkworm Bombyx mori L. using a piggyBac transposon-derived vector. Nature Biotechnology, 18 (1), 81–84.

Truett, G.E., Heeger, P.S., Mynatt, R.L., Truett, A.A., Walker, J.A., and Warman, M.L., 2000. Preparation of PCR-quality mouse genomic DNA with hot sodium hydroxide and tris (HotSHOT). Biotechniques, 29 (1), 52–54.

Wang, M., Lin, Y., Zhou, S., Cui, Y., Feng, Q., Yan, W., and Xiang, H., 2020. Genetic mapping of climbing and mimicry: two behavioral traits degraded during silkworm domestication. Frontiers in Genetics, 11, 1568.

Wang, Y., Huang, Y., Xu, X., Liu, Z., Li, J., Zhan, X., Yang, G., You, M., and You, S., 2020. CRISPR/Cas9-based functional analysis of yellow gene in the diamondback moth, Plutella xylostella. Insect Science, 28 (5), 1504–1509.

Wang, Y., Li, Z., Xu, J., Zeng, B., Ling, L., You, L., Chen, Y., Huang, Y., and Tan, A., 2013. The CRISPR/Cas System mediates efficient genome engineering in Bombyx mori. Cell Research, 23 (12), 1414–1416.

Wilkins, A.S., Wrangham, R.W., and Tecumseh Fitch, W., 2014. The “domestication syndrome” in mammals: a unified explanation based on neural crest cell behavior and genetics. Genetics, 197 (3), 795–808.

Xiang, H., Liu, X., Li, M., Zhu, Y., Wang, L., Cui, Y., Liu, L., Fang, G., Qian, H., Xu, A., Wang, W., and Zhan, S., 2018. The evolutionary road from wild moth to domestic silkworm. Nature Ecology & Evolution, 2 (8), 1268–1279.

Yamaguchi, J., Mizoguchi, T., and Fujiwara, H., 2011. siRNAs induce efficient RNAi response in Bombyx mori embryos. PLoS ONE, 6 (9), e25469.

Yoda, S., Yamaguchi, J., Mita, K., Yamamoto, K., Banno, Y., Ando, T., Daimon, T., and Fujiwara, H., 2014. The transcription factor Apontic-like controls diverse colouration pattern in caterpillars. Nature Communications, 5 (1), 4936.

Yu, H.-S., Shen, Y.-H., Yuan, G.-X., Hu, Y.-G., Xu, H.-E., Xiang, Z.-H., and Zhang, Z., 2011. Evidence of selection at melanin synthesis pathway loci during silkworm domestication. Molecular Biology and Evolution, 28 (6), 1785–1799.

Zhang, L., Martin, A., Perry, M.W., van der Burg, K.R.L., Matsuoka, Y., Monteiro, A., and Reed, R.D., 2017. Genetic basis of melanin pigmentation in butterfly wings. Genetics, 205 (4), 1537–1550.

Zhu, Y.N., Wang, L.Z., Li, C.C., Cui, Y., Wang, M., Lin, Y.J., Zhao, R.P., Wang, W., and Xiang, H., 2019. Artificial selection on storage protein 1 possibly contributes to increase of hatchability during silkworm domestication. PLOS Genetics, 15 (1), e1007616.

